# Adenine Base Editing Potently Suppresses Hepatitis B Surface Antigen Expression and Inhibits Hepatitis D Virus Release

**DOI:** 10.64898/2026.02.06.704371

**Authors:** Anuj Kumar, Emmanuel Combe, Elena M Smekalova, Selam Dejene, Dominique Leboeuf, Chao-Ying Chen, Léa Mougené, Marine Deleume, Caroline Scholtes, Marie-Laure Plissonnier, Xavier Grand, Maria G Martinez, Giuseppe Ciaramella, Francine Gregoire, Michael S. Packer, Barbara Testoni, Fabien Zoulim

## Abstract

**Background and Aims:** Novel antiviral approaches capable of permanently inactivating the intrahepatic HBV DNA reservoir, the covalently closed circular DNA (cccDNA) and HBV DNA integrated into the host genome, are urgently needed. This study evaluated adenine base editing as a strategy to disrupt HBV replication by introducing mutations in the overlapping HBs/polymerase open reading frame (ORF).

**Methods:** An adenine base editor (ABE) and 3 guide RNAs (gS1-gS3) were designed to introduce missense mutations within the HBs/polymerase ORF. ABE mRNA and individual gRNAs were co-transfected into HBV-infected HepG2-hNTCP cells and primary human hepatocytes. Antiviral efficacy was further assessed in HepG2.2.15 and PLC/PRF/5 cells harboring integrated HBV DNA. In vivo, lipid nanoparticles (LNP)-mediated delivery of ABE mRNA and gRNAs was evaluated in HBVcircle DNA-transduced mice and in HBV-infected human liver-chimeric mice. The impact of HBs editing on hepatitis D virus (HDV) release was assessed using PLC/PRF/5 and Huh7 cell-based HDV replication models.

**Results:** Adenine base editing efficiently reduced HBsAg production and HBV replication in vitro by targeting both cccDNA and integrated HBV DNA. A single LNP injection of ABE-gS2 resulted in undetectable HBsAg in HBVcircle mice, while two injections achieved a 90% reduction in serum HBsAg in HBV-infected human liver chimeric mice. HBV DNA replication was also inhibited in vivo. Furthermore, HBs ORF base editing markedly suppressed HDV release in vitro.

**Conclusions:** Adenine base editing of the HBs ORF effectively impairs HBV replication and HBsAg production in vitro and in vivo and concomitantly inhibits HDV release, highlighting its therapeutic potential.

## Introduction

With over 250 million chronic infections and approximately 1 million deaths per year globally, Hepatitis B virus (HBV) represents a serious health burden (1). Currently approved nucleos(t)ide analogues (NAs) efficiently suppress HBV replication by inhibiting viral reverse transcription. However, they require lifelong treatment for patients with chronic hepatitis B (CHB). Importantly, NAs do not directly target the HBV genomic reservoir which includes the viral episomal minichromosome (cccDNA) and HBV DNA integrated in the host genome. In addition to HBV DNA undetectability, hepatitis B surface antigen (HBsAg) in serum is considered the optimal endpoint to achieve a functional cure for CHB (2–5). HBsAg is also central in the replication cycle of the satellite virus HDV, that requires HBs proteins for its envelopment and release from infected cells to establish a chronic co-infection.

Emerging treatments such as small interfering RNA (siRNA), antisense oligonucleotides (ASOs), and Cas13b, target the HBV viral replication cycle downstream of these key viral genome species (6–10), and strategies aimed at inhibiting HBsAg secretion/production are being explored to prevent HDV dissemination (11).

To achieve permanent cure, several research groups have explored CRISPR/Cas9 nuclease-or homing endonuclease (ARCUS nuclease)- based strategies to eliminate or inactivate cccDNA and integrated HBV DNA by introducing double strand breaks (DSBs) sensitive to cellular nucleases and/or by introducing mutations or deletions following DNA repair (12–14). However, this approach is associated with significant challenges, such as the risk of undesired consequences including insertions, deletions and chromosomal rearrangements, that may lead to genomic instability (15–18).

Recent advances in the CRISPR/Cas9 field have led to the development of gene editing tools that do not require DSB-intermediates such as base editors, which have transformed the gene editing field (19–21). At least four independent studies have shown the potential of cytosine base editors (CBEs) to functionally inactivate HBV genomes by introducing C-to-T substitutions, leading to the introduction of non-sense/missense mutations in HBV DNA species (22–25). Although prototypical CBEs, containing naturally occurring cytidine deaminase, are less likely to cause genomic rearrangements than nucleases (26), they can still be associated with off-target effects including guide-independent cytosine deamination of cellular DNA and RNA (27–29). Next-generation CBEs can mitigate such off-target effects; however, these advanced base editors may exhibit reduced on-target efficiency (30). In contrast, adenine base editors (ABEs), which utilize a laboratory-evolved TadA adenine deaminase to convert A-to-G, demonstrated high on-target and minimal off-target editing, making them preferred for precision gene-editing (31,32). Furthermore, studies have confirmed the therapeutic efficacy of ABEs in vivo (33–35). To date, only one study demonstrated the inhibition of HBV replication and antigens expression by plasmid-based delivery of ABEs (25). In the present study, we demonstrate the efficacy of mRNA-based delivery of ABEs in inhibiting HBV replication and HBsAg expression from both cccDNA and integrated HBV DNA using de novo infected HepG2-hNTCP, primary human hepatocytes (PHHs), as well as HepG2.2.15 and PLC/PRF/5 stable cell lines. Furthermore, HBsAg loss or sustained HBsAg decrease was observed in vivo in HBVcircle and HBV-infected human liver chimeric mouse models, respectively. Additionally, we provide in vitro ‘proof-of-concept’ that targeting HBsAg can be used as a strategy to inhibit HDV release and, hence, its spread. Altogether, adenine base editing represents a promising approach for potent suppression of HBsAg expression and consequently inhibiting HDV release.

## Materials and methods

### 2.1 Base editing reagents

Base editing was performed by utilizing mRNA encoding an 8^th^ generation adenine base editor (31,32), and HBs/POL-targeting gRNAs named gS1, gS2, and gS3. A gRNA targeting the host *ALAS1* gene was used as a non-HBV targeting control gRNA (36).

### 2.2 Plasmids

The plasmid pSVLD3 containing head-to-tail trimer of full length HDV cDNA assembled into the SV40 based vector pSVL (37,38) was used to produce HDV particles in PLC/PRF/5 and in Huh7 cells. The plasmid pT7HB2.7 containing preS1-preS2-S gene sequence was used to express L, M and S isoforms of HBsAg in Huh7 cells(39). HBs ORF from genotypes A-H were generated from a consensus analysis of the HBVdb database (40), subcloned by Genscript and aligned to clinical isolates sequences OP611181 (GT-A), KJ173408 (GT-B), KJ173426 (GT-C), U95551 (GT-D), GQ161816 (GT-E), KP995098 (GT-F), KY004111 (GT-G) and AB375162 (GT-H).

### 2.3 Cell culture experiments

Cell lines. HepG2-hNTCP, HepG2.2.15 cells and PLC/PRF/5 cells were cultured as previously described (24). Huh7 cells were maintained in DMEM supplemented with 10% fetal calf serum (Fetal clone II), 1% penicillin-streptomycin (Life Technologies), 1% GlutaMAX (Gibco), and 1% sodium pyruvate (Gibco). For HDV production in Huh7 cells, complete William’s medium was used. Twenty four-well plates of primary human hepatocytes (PHHs) isolated from chimeric mouse liver were purchased from Phoenix Bio and maintained in defined hepatocyte growth medium (dHCGM).

#### Production of HBV for infection experiments in vitro

HBV (genotype D, subtype ayw) infectious Dane particles were generated by polyethylene-glycol-MW-8000 (PEG8000, SIGMA) precipitation of the supernatants obtained from HepAD38 cells as previously described (41).

#### Adenine base editing in HepG2-hNTCP

HepG2-hNTCP cells were trypsinized and plated in complete DMEM at a density of 10^5^ cells/cm^2^. Starting the following day, the cells were cultured in medium containing 2.5% DMSO (Merck-Sigma-Aldrich) to promote differentiation and enhance HBV infection. After 72 hours, the cells were infected with HBV at a multiplicity of infection (MOI) of 1000. The infected cells were replated on day 6 post-infection (6 dpi). On the following day (i.e. 7 dpi), the cells were transfected with ABE-encoding mRNA and gRNA at a ratio of 2:1 (200ng:100ng) using Lipofectamine Messenger MAX (Life Technologies) and Opti-MEM (Gibco). At 15dpi, supernatants and cells were harvested to assess extracellular and intracellular HBV parameters, respectively.

For experiments involving Lamivudine (LAM) treatment (**Fig. S1**), 10 µM LAM was added at 4 dpi. The cells were replated at 6dpi, and LAM treatment was maintained until the final day of the experiment (15dpi).

#### Adenine Base editing in PHHs

PHHs were infected with HBV at an MOI of 500. At 4 dpi, after the establishment of a stable pool of cccDNA, base editing was performed. PHHs in 24-well plates were treated with 1 µg/well of total RNA formulated in lipid nanoparticles. At 18 dpi, supernatants were collected to evaluate extracellular viral parameters, and cells were harvested to analyze intracellular HBV parameters and base editing efficiency.

#### Adenine base editing in PLC/PRF/5 and HepG2.2.15 cells

PLC/PRF/5 cells were transfected with ABE-encoding mRNA (200ng) and gRNA (100ng). At 3 days post-transfection, extracellular HBsAg levels in the supernatants, and HBs ORF editing efficiency from cell lysates were analyzed. For experiments with HepG2.2.15 cells, these cells were pre-treated with LAM before transfection with ABE-encoding mRNA (200ng) and gRNA (100ng). At 7 days post-transfection, different HBV parameters were analyzed.

### 2.4 In vivo evaluation of ABE in mice

#### Ethical statement for in vivo experiments

HBVcircle mice experiments were performed according to the relevant National Institutes of Health guidelines and were approved by the Institutional Animal Care and Use Committee and the Office of Laboratory Animal Research at Charles River Accelerator and Development Laboratory. The PXB mouse study was governed by the Animal Ethics Committee of PhoenixBio.

#### Lipid Nanoparticles (LNPs) formulations

The base editor mRNA and gRNA were co-encapsulated at a 1:1 weight ratio in LNPs. The LNPs were generated by rapidly mixing an aqueous solution of the RNA at a pH of 4.0 with an ethanol solution containing four lipid components: an ionizable lipid, distearoylphosphatidylcholine (DSPC), cholesterol, and a lipid-anchored polyethylene glycol (PEG) (all of the lipids were obtained from Avanti Polar Lipids). The two solutions were mixed using the benchtop microfluidics device from Precision Nanosystems. Post-mixing, the formulations were dialyzed overnight at 4°C against 1× Tris-buffered saline (TBS) (Millipore Sigma). They were subsequently concentrated down using 100,000 Da molecular weight cutoff (MWCO) Amicon Ultra centrifugation tubes (Millipore Sigma) and filtered with 0.2-μm filters (Pall Corporation). The total RNA concentration was determined using Quant-iT Ribogreen (Thermo Fisher Scientific).

#### HBVcircle experiment

Immunocompetent C3H male mice, aged 5–6 weeks, were obtained from Taconic Biosciences. Over the course of the study mice were group housed (up to 5 animals together) in individually ventilated, solid bottom caging with Alpha-Dri® contact bedding material, given enrichment items per site standard procedures (igloo, Nestlets or nesting strips), and fed Prolab Isopro RMH 3000 (LabDiet). All interventions and sample collections were performed in the unfasted state during the light cycle. HBVcircle was delivered into these mice through hydrodynamic injection (HDI) as previously described (24,42). 28 days post-HDI, mice were screened for HBsAg levels and subsequently grouped into four groups as shown in Fig.4. LNPs were administered at 2mg/kg (total RNA/body weight) or 0.3mg/kg entecavir was administered intravenously through the tail vein. Sera were collected via the submandibular vein to monitor HBsAg levels. The animals were euthanized 33 days after LNP injection.

#### HBV infected human liver chimeric mice experiment

Male PXB human liver chimeric mice aged 23-26 weeks were infected with HBV (genotype C). Over the course of the study mice were individually housed in TP-112 caging (TOYO-LABO) with sterilized soft paper bedding (PaperClean), given Diamond Twist enrichments (Envigo), and fed CRF1 chow and AS supplement (Oriental Yeast Co). All interventions and sample collections were performed in the unfasted state during the light cycle. Mice were assigned to groups to achieve similar average body weight, blood human albumin, and serum HBV DNA. LNPs were administered two times at 2mg/kg (total RNA/body weight) intravenously through the tail vein. Sera were collected via the retro-orbital plexus/sinus to analyze HBsAg levels. The animals were euthanized 56 days after first LNP injection. These experiments were performed at Phoenix Bio Co; Ltd. (Hiroshima, Japan).

### 2.5 Analysis of viral proteins

#### ELISA to quantify extracellular HBsAg

HBsAg levels in cell supernatants of infected HepG2-hNTCP, HepG2.2.15 and PLC/PRF/5 or in the mice sera for the HBVcircle experiment were measured by ELISA using a chemiluminescence immunoassay (CLIA, lower limit of detection = 0.04 IU/ml) kit (Autobio Diagnostic, Zhengzhou, Henan, CN), according to the manufacturer’s protocol. HBsAg levels in supernatants of infected PHHs was measured by HBsAg ELISA GS HBsAg EIA 3.0, Bio-Rad Laboratories, Inc., Los Angeles CA, Supplier Part# 32591). HBsAg levels in the mice sera for the PXB mouse experiment were quantified by ELISA developed by Fujirebio (LUMIPULSE HBsAg-HQ, LUMIPULSE® L2400).

#### Western blot

40µg of lysates were resolved on 4-20% or 12% mini-PROTEAN TGX stain-FreeTM Precast gels (Bio-Rad Laboratories) and transferred onto nitrocellulose membranes (Bio-Rad Laboratories). Immunodetection was performed using primary antibodies including monoclonal anti-HBs (Abbott H166 mouse), polyclonal anti-HBs (Novus biologicals NB100-62652 rabbit), anti-Ku80 (ab119935, mouse, Abcam), anti-HDV (homemade, mouse), followed by incubation with horseradish peroxidase (HRP) or fluorophore-conjugated secondary antibodies. Blots were visualized using Clarity Western ECL (Bio-Rad Laboratories) and the Chemidoc MP Imaging system (Bio-Rad Laboratories).

### 2.6 Quantification of intracellular HBV DNA

Total intracellular DNA was extracted using the MasterPure Complete DNA and RNA Purification Kit (Lucigen). HBV DNA was quantified via qPCR using the TaqMan assay Pa03453406_s1(Life Technologies). For cccDNA detection, total intracellular DNA was digested with exonucleases I and III before qPCR quantification using the primers Forward: 5’CCGTGTGCACTTCGCTTCA30; Reverse: 50GCACAGCTTGGAGGCTTGA30; and the probe: [6FAM] CATGGAGACCACCGTGAACGCCC [BBQ]. qPCR was performed using an Applied QuantStudio 7 real time machine. Hemoglobin subunit beta (HBB, TaqMan Assay ID: Hs00758889_s1) housekeeping gene served as an internal control for normalization.

### 2.7 Assessment of HBs-ORF editing

Extracted DNA samples were analyzed via targeted amplicon sequencing. Briefly, sample preparation includes target-specific amplification with Q5 Hot Start High-Fidelity DNA Polymerase, barcoding reactions with unique Illumina barcoding primer pairs, amplicon purification, and DNA quantification. The library was diluted and assessed on the next-generation sequencing Illumina MiSeq platform. Editing rates were determined by analyzing sequencing reads as described previously (43) and with the CRISPResso2 software pipeline (44).

### 2.8 Analysis of HDV

#### HDV production in PLC/PRF/5 cells

PLC/PRF/5 cells were seeded at a density of 0.5 x10^5^ cells/cm^2^. After 24 hours, transfection with pSVLD3 plasmid was performed using TransIT®-2020 Transfection Reagent (Mirus Bio) and Opti-MEM following the manufacturer’s instructions. The following day, cells were washed 3 times with 1X PBS and maintained in complete MEM medium. Supernatants and cells were collected to assess different viral parameters.

#### HDV production in Huh7 cells

HuH7 were seeded at a density of 1×10^5^ cells/cm^2^ in William’s medium. After 24 hours, cells were co-transfected with pSVLD3 and pT7HB2.7 plasmids (ratio 1:1) using TransIT®-2020 Transfection Reagent (Mirus Bio) according to the manufacturer’s instructions. The next day, cells were washed 3 times with 1X PBS and maintained in William’s medium. Supernatants and cells were harvested to assess different viral parameters.

#### Quantification of extracellular HDV RNA

A fixed volume of 200 µl of HDV producing supernatant was used to extract viral nucleic acid using the High Pure Viral Nucleic Acid Kit (Roche LifeScience) following the manufacturer’s instructions. Residual expression plasmids were removed by performing RQ1 RNase-Free DNase step (Promega) prior to HDV RNA quantification. HDV RNA levels were measured by one-step RT-qPCR using the Fast Virus 1-Step TaqMan™ Master Mix (Life Technologies SAS) and an HDV probe [6FAM]-ATGCCCAGGTCGGAC-[BHQ®-1] along with primers (Forward 5’-TGGACGTGCGTCCTCCT-3’; Reverse 5’-TCTTCGGGTCGGCATGG-3’) (45). An *in vitro* transcribed HDV RNA was used as an external standard to monitor the efficiency of RT-qPCR reactions.

## Results

### 3.1 Adenine base editing efficiently suppresses HBsAg and inhibits HBV replication in infected hepatocytes in vitro

We investigated the effect of ABE with different gRNAs on the production of HBsAg from established cccDNA in HepG2-hNTCP cells, following a protocol similar to that used previously for cytosine base editing (24) (**Fig.1A**). HBV-infected HepG2-hNTCP cells were co-transfected with ABE-expressing mRNA and individual gRNAs on day 7^th^ post infection. On the 8th day post-transfection (i.e., day 15^th^ post infection), secreted HBsAg levels were measured by ELISA. Profound reductions of 85%, 90% and 79% in HBsAg levels were observed with gS1, gS2 and gS3, respectively (**Fig.1B**). Due to the overlapping nature of HBV open reading frames (ORFs), base editing of the HBs ORF at specific target sites by these gRNAs also results in concomitant missense mutations in the RT domain of POL ORF, potentially affecting HBV replication. Therefore, we also examined the effect of adenine base editing of HBs/POL ORF by all three gRNAs on HBV replication by quantifying total intracellular HBV DNA levels in vitro in HBV-infected HepG2-hNTCP. Interestingly, gS1 and gS3 demonstrated a robust reduction, while only a partial reduction was observed with gS2 (**Fig. 1C**), without impacting cccDNA levels (**Fig. 1D**). On-target editing efficiency of 32 to 54 % of HBs ORF on cccDNA was obtained for the HBV- targeting gRNAs (**Fig. 1E**).

**Fig. 1.**
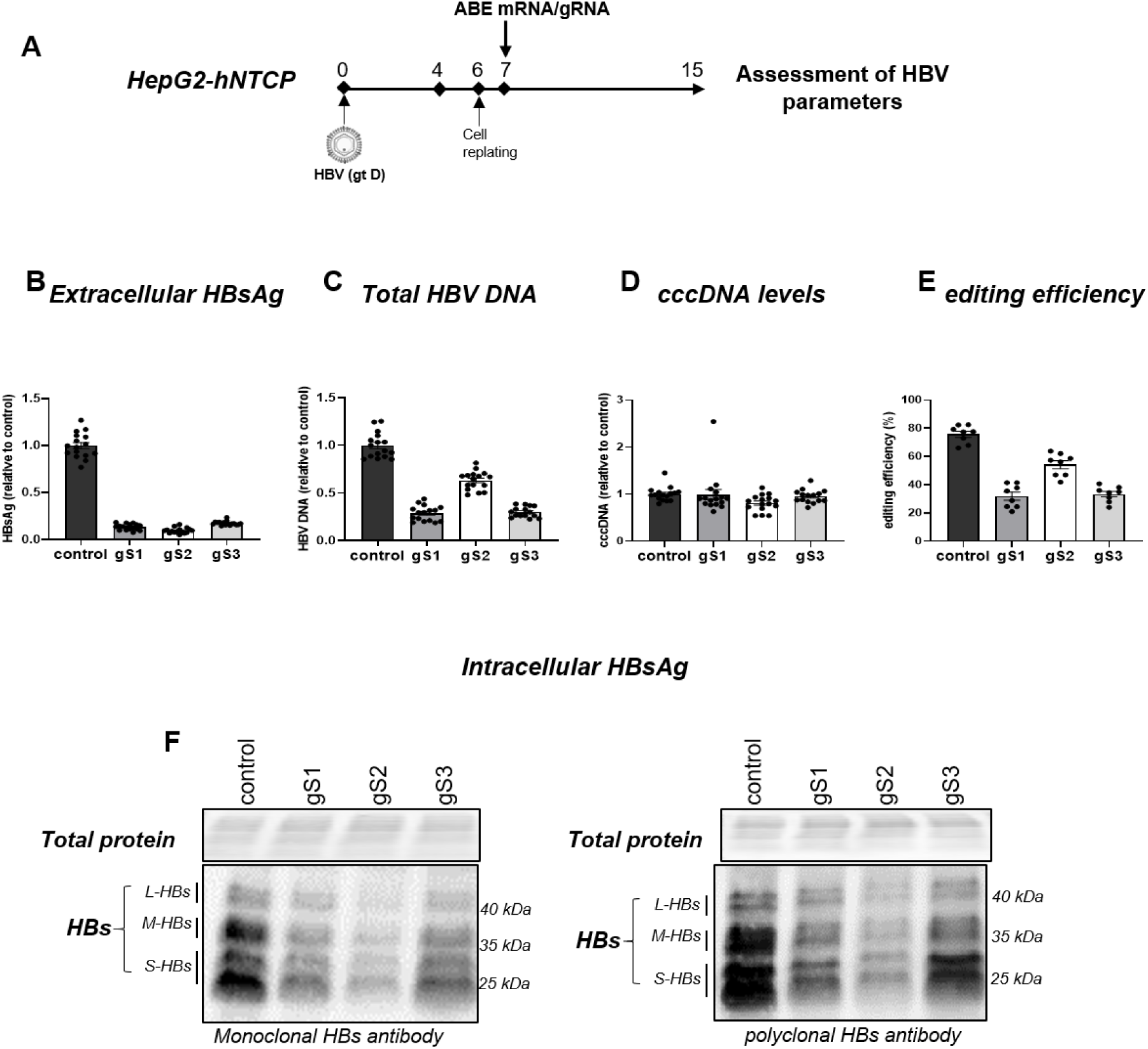
ABE/gRNAs drastically reduce extracellular HBsAg and inhibit HBV DNA in vitro in hepatoma cells. (A) Schematic of the protocol used to transfect HBV-infected HepG2-hNTCP cells with ABE-encoding mRNA and HBs/POL ORF targeting gRNA (gS1, gS2, or gS3). (B) The levels of extracellular HBsAg were determined by ELISA. (C) intracellular total HBV DNA and (D) cccDNA levels were quantified by qPCR. (E) Percentage of A-to-G editing of HBs ORF by gS1-gS3 and control gRNA was detected on exonucleases I/III-treated samples and total extracted DNA samples, respectively. control: non-HBV targeting gRNA. Data are represented as mean±SEM. (F) The levels of intracellular HBsAg were determined by Western blot using anti-HBs monoclonal (H166, Abbot) or polyclonal antibody (NB100-62652, NovusBio). Ponceau staining of total protein was used to control protein loading.

Consistent with ELISA results, Western blotting using monoclonal HBs antibody H166 demonstrated reduced intracellular HBsAg levels, with gS2 showing the highest efficacy, followed by gS1 and then gS3 (**Fig. 1F**). Similar results were obtained using a polyclonal antibody against HBs, suggesting that the decreased intracellular HBsAg levels would most probably be associated with ABE-induced HBs missense mutations rather than with an altered antigen detection derived from amino acid changes (**Fig. 1F**). Importantly, the effect of adenine base editing on extracellular and intracellular parameters was maintained in the presence of LAM treatment (**Fig. S1, A-E)**.

Next, we assessed the ability of the most effective HBs reducing gRNAs, gS1 and gS2, to decrease HBsAg expression in infected PHHs. A single treatment with LNPs formulated with ABE-encoding mRNA and HBs ORF-targeting gRNA was performed. PHHs were kept in culture for 14 more days, with the experiment concluding 18 days post-infection (**Fig. 2A**). Consistent with the HepG2-hNTCP data, adenine base editing resulted in a robust reduction of greater than 90% in extracellular HBsAg by gS1 and gS2 (**Fig. 2B**). Similarly, for HBV DNA replication, gS1 demonstrated a robust reduction, while only a partial reduction was observed with gS2 (**Fig. 2C**). NGS-based amplicon sequencing confirmed the robust editing of HBs ORF (**Fig. 2D**). Altogether, these findings suggest that adenine base editing can effectively suppress HBsAg levels and reduce HBV replication in vitro.

**Fig. 2.**
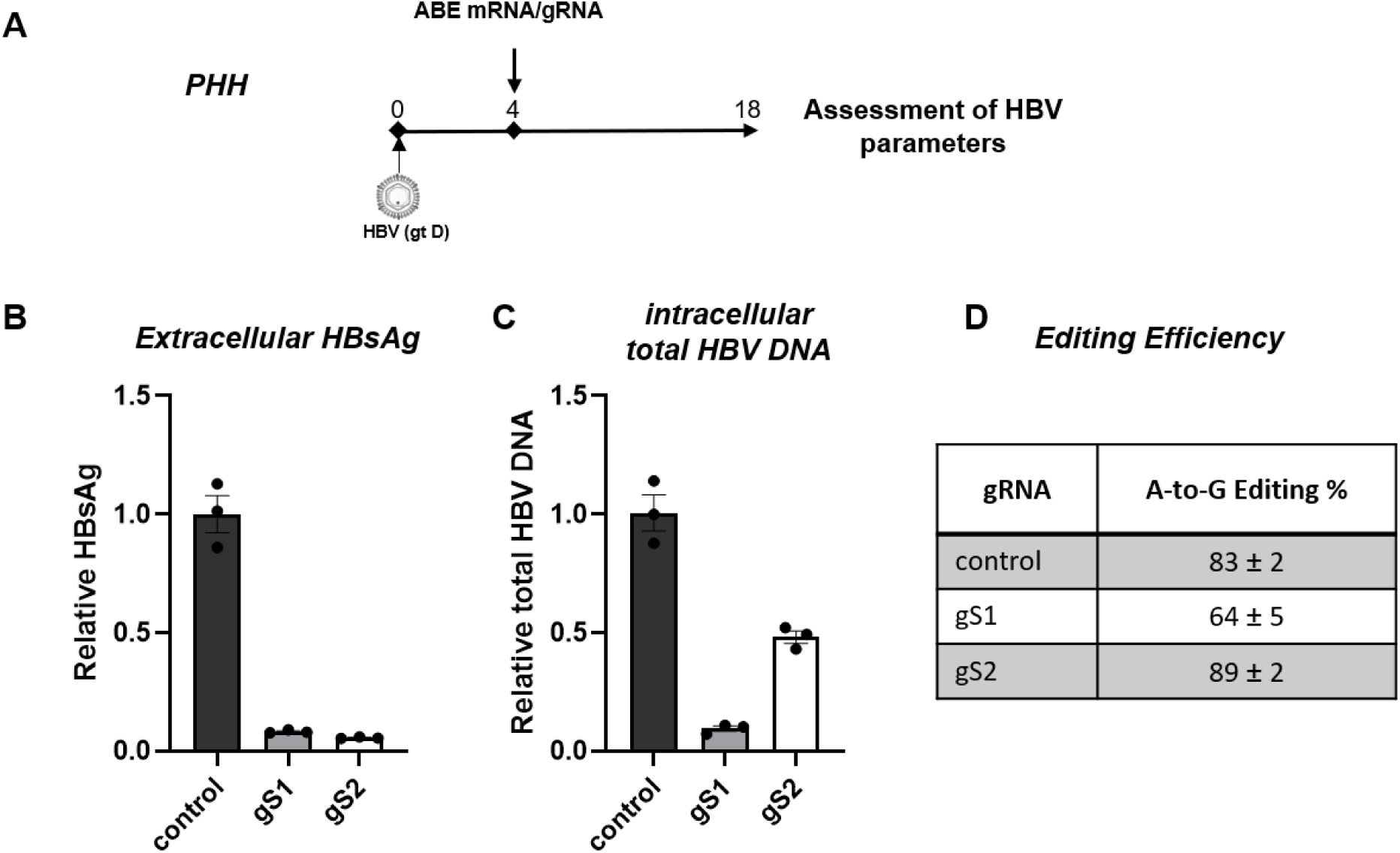
ABE/gRNAs drastically reduce extracellular HBsAg and inhibit HBV DNA in vitro in primary human hepatocytes. (A) Schematic of the protocol used to treat HBV-infected PHHs with ABE-encoding mRNA and gS1 or gS2. (B) The levels of extracellular HBsAg and (C) intracellular total HBV DNA were determined by ELISA and qPCR, respectively. (D) A-to-G editing % of HBs ORF (for gS1, gS2, and gS3) and *control* gene (for control gRNA) was determined by NGS-based amplicon sequencing. control: non-HBV targeting gRNA. Data are represented as mean ± SEM.

### 3.2 Adenine base editing decreases HBsAg derived from integrated HBV DNA

As integrated HBV DNA represents an important source of HBsAg, especially in HBeAg-negative patients (46), we next investigated whether gS1 and gS2 could inhibit HBsAg expression from integrated HBV genomes. To address this, we utilized HepG2.2.15 cells, which harbor engineered integrated HBV DNA in host cell genome. These cells were treated with LAM prior to transfection with ABE-encoding mRNA and gS1 or gS2 (**Fig. 3A**). A robust reduction of extracellular HBsAg (**Fig. 3B**), as well as intracellular HBsAg **(Fig. S2)** was observed, with 76% and 77% editing of the integrated HBs ORF with gS1 and gS2, respectively (**Fig. 3C**).

**Fig. 3.**
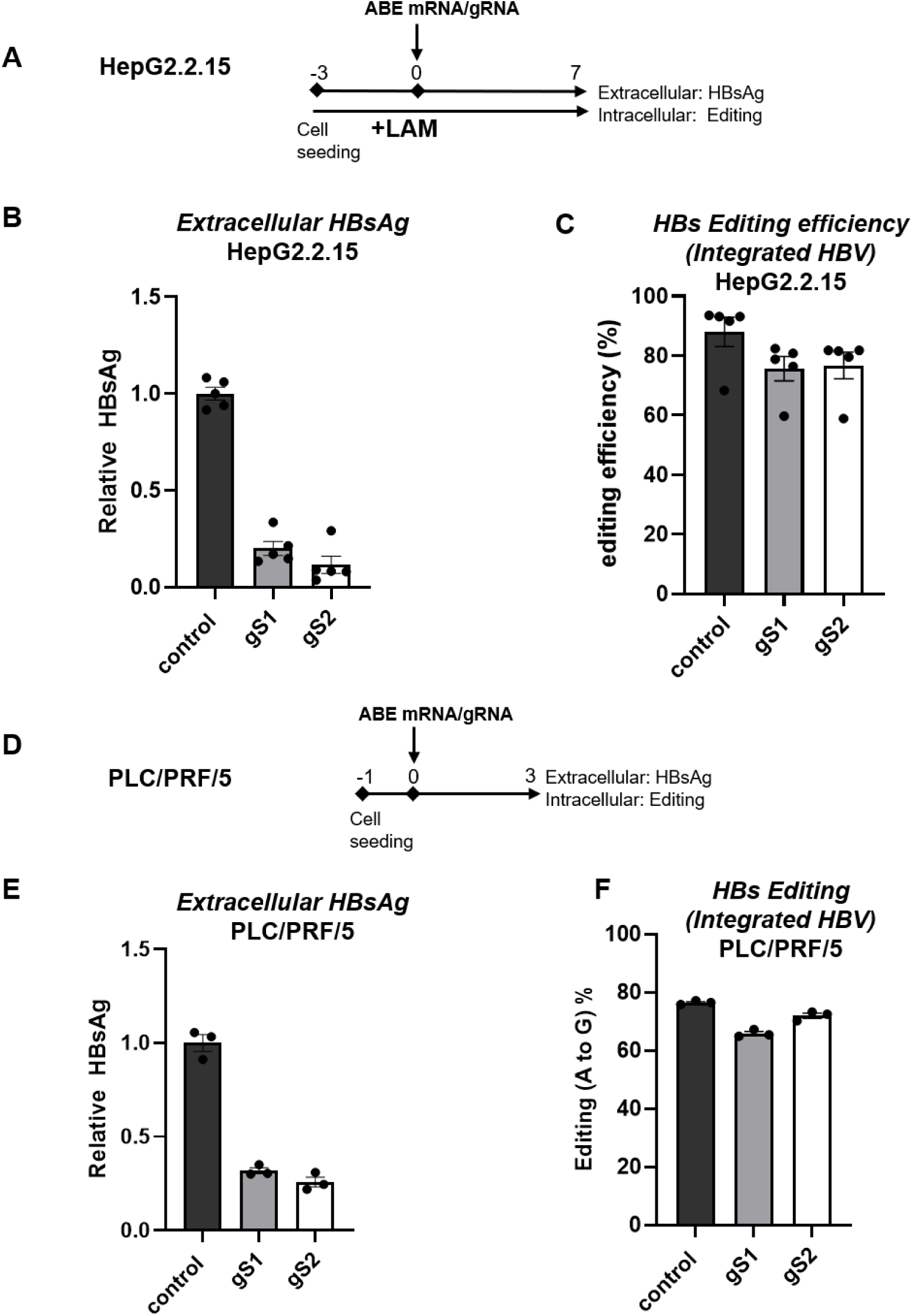
ABE/gRNAs drastically reduce extracellular HBsAg from integrated HBV DNA. (A). The schematic of the protocol used to transfect LAM pretreated HepG2.2.15 cells with ABE-encoding mRNA and gS1 or gS2. (B) The levels of extracellular HBsAg and (C) the percentage of HBs ORF editing in HepG2.2.15 cells were determined by NGS-based amplicon sequencing. (D) The schematic of the protocol used to transfect PLC/PRF/5 with ABE-encoding mRNA and gS1 or gS2. (E) The levels of extracellular HBsAg and (F) the percentage of HBs ORF editing in PLC/PRF/5 cells. control: non-HBV targeting gRNA; LAM: Lamivudine. Data are represented as mean ± SEM.

**Fig. 4.**
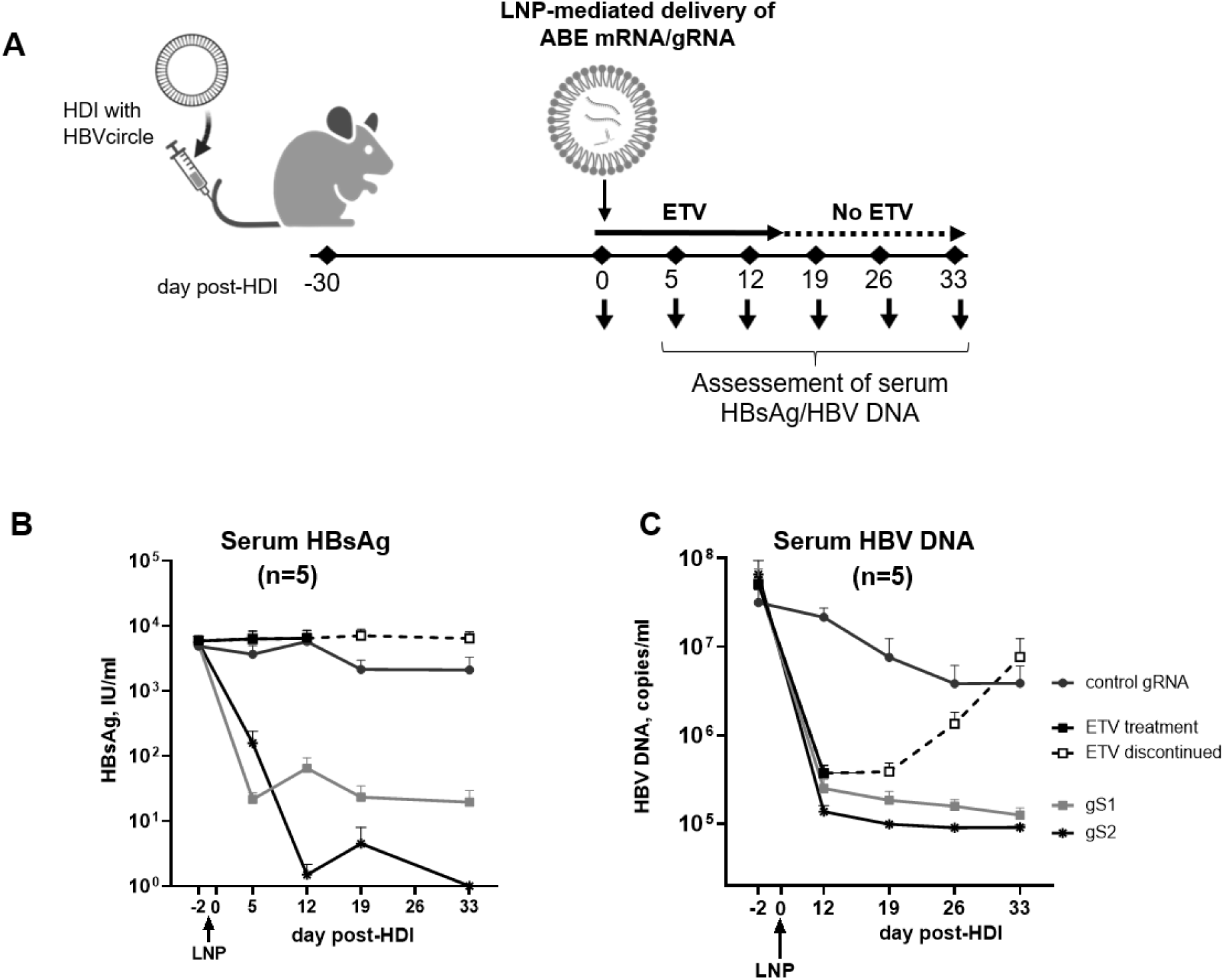
Adenine base editing leads to sustained reduction of HBsAg and HBV DNA in HBVcircle mouse model. (A) Schematic of the protocol used to administer LNPs encapsulating ABE-encoding mRNA and HBs-targeting gRNA (gS1or gS2) followed by assessment of serum HBsAg and HBV DNA in the HBVcircle mouse model. (B) Serum HBsAg levels were determined by ELISA at different days post first LNP injection. (C) In vivo efficacy of gS1 and gS2 on serum HBV DNA levels. control: non-HBV targeting gRNA; ETV: entecavir. Data are represented as mean ± SEM. Partially created in BioRender. Testoni, B. (2026) https://BioRender.com/cnmc79r

The effect of adenine base editing on HBs expression was further analyzed in PLC/PRF/5 cells, that harbor replication incompetent naturally integrated HBV DNA sequences (genotype A) (**Fig. 3D**). Similar to the results obtained with HepG2.2.15 cells, around 70 and 75% reduction of extracellular HBsAg was observed with gS1 and gS2, respectively (**Fig. 3E**). 65-72% of HBs ORF editing was sufficient to achieve the observed HBsAg decrease in this cellular model (**Fig. 3F**).

### 3.3 Pangenotypic impact of editing mutations on HBs levels

As gS2 demonstrated the most efficient suppression of HBsAg levels from HBV gt D (HepG2-hNTCP, PHH, HepG2.2.15) and HBV gt A (PLC/PRF/5), we next investigated the impact of ABE gS2 induced mutations on HBsAg across eight HBV genotypes (A to H). These mutations were engineered into the pT7HB2.7 plasmid containing the complete HBs nucleotide sequence derived from genotypes A to H. Huh7 cells were transiently transfected with either WT HBV or ABE gS2-mutant plasmid. At six days post-transfection, secreted and intracellular HBsAg levels were determined in culture supernatants and cell lysates, respectively (**Fig. S3A)**.

A marked suppression in HBsAg levels was observed in the ABE mutants across all HBV genotypes when compared to their respective wildtype (WT) controls **(Fig. S3B-S3C)**. qPCR analysis of the transfected plasmids revealed no difference in transfection efficiency between WT and mutant constructs (data not shown), suggesting that these mutations indeed lead to drastic and pangenotypic decrease of HBsAg.

### 3.4 Adenine base editing leads to sustained inhibition of HBsAg in mouse models

To examine the impact of adenine base editing on HBsAg expression in vivo, we first employed the HBVcircle mouse model (42). Mice were hydrodynamically injected with HBVcircle, an HBV genotype D cccDNA-like plasmid. Thirty days post-injection, the mice received a single systemic administration of lipid nanoparticles (LNPs) encapsulating ABE-encoding mRNA and HBs-targeting gRNA (gS1 or gS2) via intravenous injection (**Fig. 4A**). Serum HBsAg was monitored at day 5, 12, 19, and 33 post-LNP injection. Interestingly, a sustained reduction of HBsAg was observed, with up to 2.5 log10 (99.68%) and 4log10 (99.99%, 5/5 mice below the lower limit of detection) suppression at day 33^rd^ post-LNP injection with gS1 and gS2, respectively (**Fig. 4B**).

We also investigated the effect of adenine base editing in the HBV-infected human liver chimeric mouse model. Mice were infected with HBV genotype C (**Fig. 5A**). When mice were in a stable phase of infection, typically >4 weeks post-infection (47), two consecutive administrations with LNPs encapsulating ABE-encoding mRNA and HBV targeting gRNAs resulted in inhibition of serum HBsAg levels at 56 dpi with up to a 90% (1log10) reduction with gRNA S2 (**Fig. 5B**). Editing efficiencies of 47% and 68% was observed in the HBs ORF with gS1 and gS2, respectively (**Fig. 5C**).

**Fig. 5.**
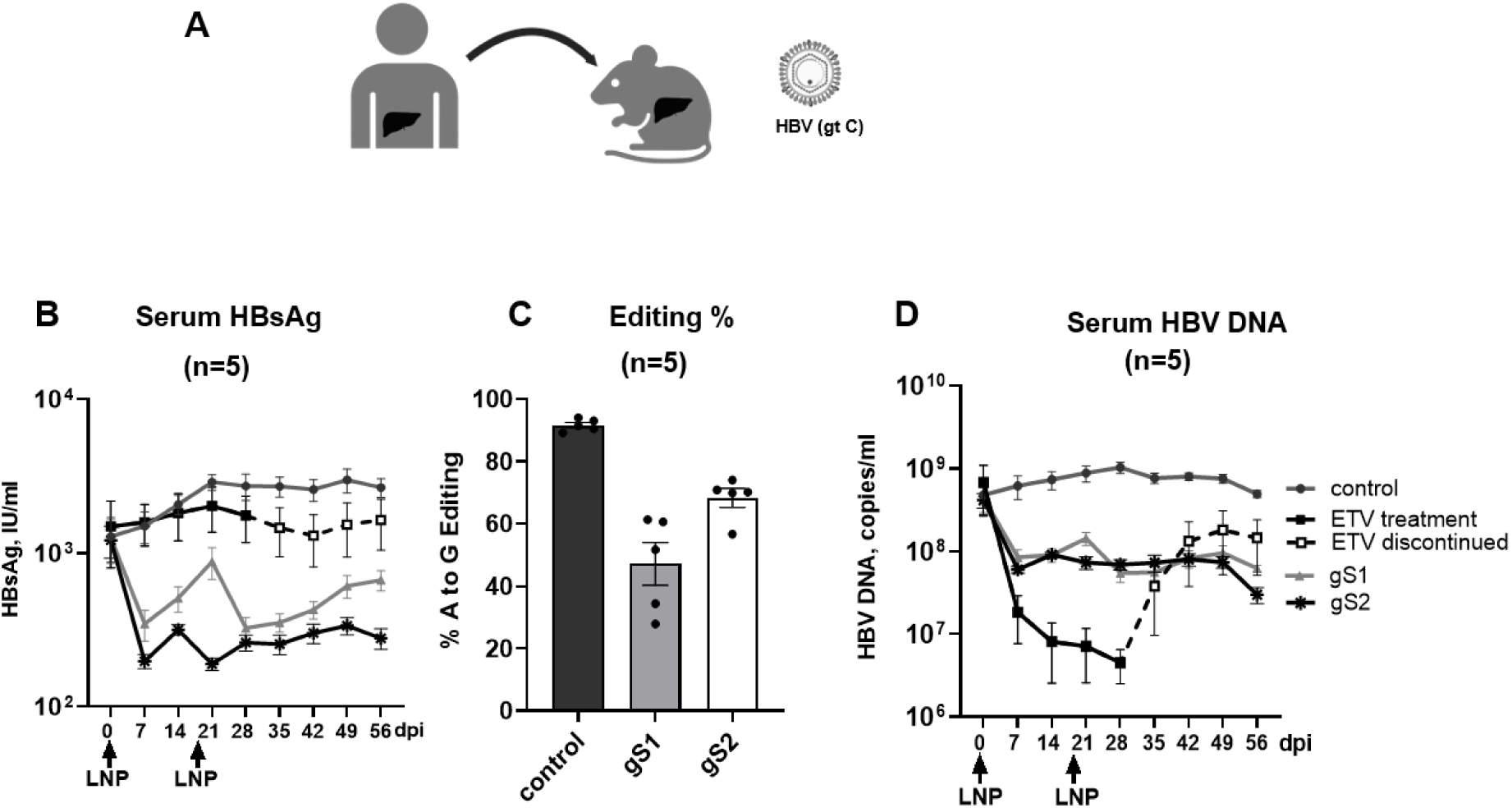
Adenine base editing leads to sustained reduction of HBsAg and HBV DNA in the HBV-infected human liver chimeric mouse model. (A) Schematic of the protocol used to administer LNPs encapsulating ABE-encoding mRNA and gS1 or gS2 followed by assessment of serum HBsAg in the HBV-infected humanized mice model. (B) Serum HBsAg levels were determined by ELISA at different days post first LNP injection. (C) HBs/POL ORF editing was determined at the end of the experiment (56 dpi). (D) In vivo efficacy of gS1 and gS2 on serum HBV DNA levels. dpi: day post first LNP injection. control: non-HBV targeting gRNA; ETV: entecavir. Data are represented as mean ± SEM. Created in BioRender. Testoni, B. (2026) https://BioRender.com/0mc27ht

As expected, in both mouse models, the animals receiving Entecavir (ETV) orally, for the first 14 days post-LNP injection in the HBVcircle model and for 28 days after the first-LNP injection in the HBV human liver chimeric mouse model, followed by treatment discontinuation (**Fig. 4B and Fig 5B**), showed no change in HBsAg levels.

Further, both gRNAs showed 1.8 log10 (98%) and at least 1log10 (90%) reduction in serum HBV DNA levels in the HBVcircle and HBV-infected human liver chimeric mice, respectively, at the end of the study (**Fig. 4C and 5D**). In contrast to the sustained reduction observed in gS1 or gS2-treated mice groups, an independent control group receiving Entecavir (ETV) orally showed HBV rebound shortly after treatment discontinuation (**Fig. 4C and 5D**).

Altogether, these results demonstrate the potential of adenine base editing to induce a sustained decrease of both HBsAg and HBV replication in vivo.

### 3.5 Adenine base editing-induced HBs mutations inhibit HDV release and spread

We investigated whether reducing HBsAg levels through adenine base editing could impact HDV release. To address this question, we used two approaches.

#### HBs ORF base Editing in PLC/PRF/5 cells

We used PLC/PRF/5 cells that are able to produce infectious HDV virions upon transfection of an HDV encoding plasmid, pSVLD3 (38). PLC/PRF/5 cells were transfected with pSVLD3, followed by transfection with ABE-encoding mRNA and gS2 to edit the HBs ORF naturally integrated in the PLC/PRF5 cell genome. Levels of HBsAg and HDV RNA in the supernatants were assessed at day 6 (d6) post base editing (**Fig. 6A**). Similar to the decrease in extracellular HBsAg levels (**Fig. 6B**), extracellular HDV RNA was found to be drastically reduced in ABE gS2 base edited cells compared to control gRNA edited cells (**Fig. 6C**). The reduction of intracellular HBsAg (**Fig. 6D**) as well as high editing (82%) of the integrated HBs sequence were also confirmed (**Fig. 6E**).

**Fig. 6.**
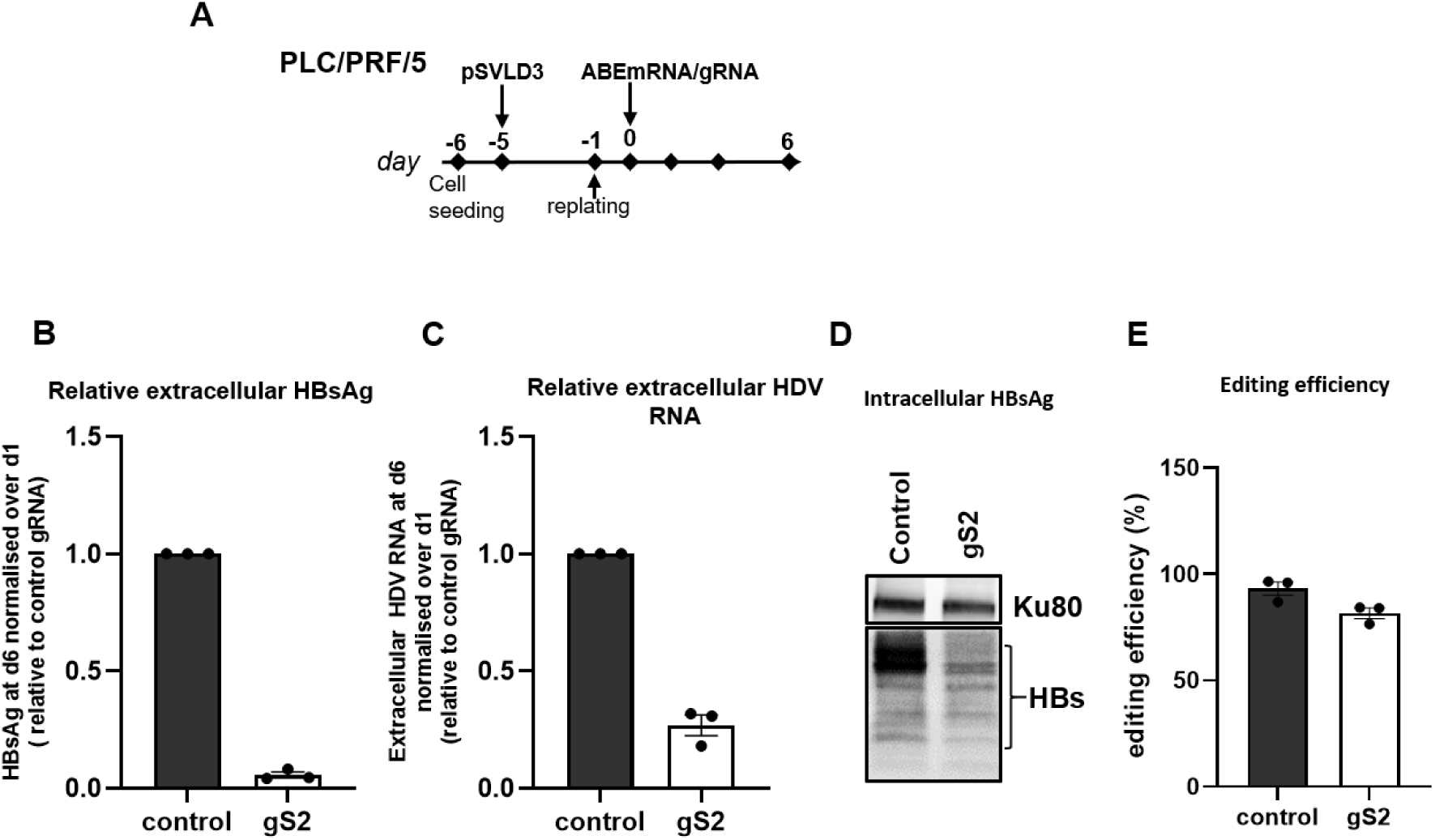
Adenine base editing reduces HDV release in PLC/PRF/5 cells. (A) Schematic of the protocol used to transfect PLC/PRF/5 cells with pSVLD3 followed by transfecting ABE-encoding mRNA and gS2. (B) Relative HBsAg (C) and extracellular HDV RNA levels were determine at day 6 (d6). (D) Western blot to detect intracellular HBsAg was performed at the end of experiment. Ku80 served as a protein loading control. (E) A-to-G editing % of integrated HBs ORF (for gS2) and *control* gene was determined at the end of experiment (for control non-HBV targeting gRNA). Data are represented as mean ± SEM.

To rule out the possibility that reduced extracellular HDV RNA observed in the gS2 condition resulted from decreased pSVLD3 transfection efficiency or intracellular HDV RNA levels, the levels of pSVLD3 plasmid and intracellular HDV RNA were quantified. No differences were observed between control gRNA and gS2-edited conditions **(Fig. S4).**

#### Effect of base editing induced HBs mutations on HDV release

To further confirm that HBs mutations led to reduced HDV RNA levels in the supernatant, we employed a complementary approach, where ABE-gS2 corresponding mutations were engineered into a plasmid, pT7HB2.7, encoding full-length HBsAg, expressing all three HBs isoforms (cf. Method section). Huh7 cells were co-transfected with pSVLD3 and pT7HB2.7 expressing wild-type (WT) or mutant HBs proteins (**Fig. 7A**).

**Fig. 7.**
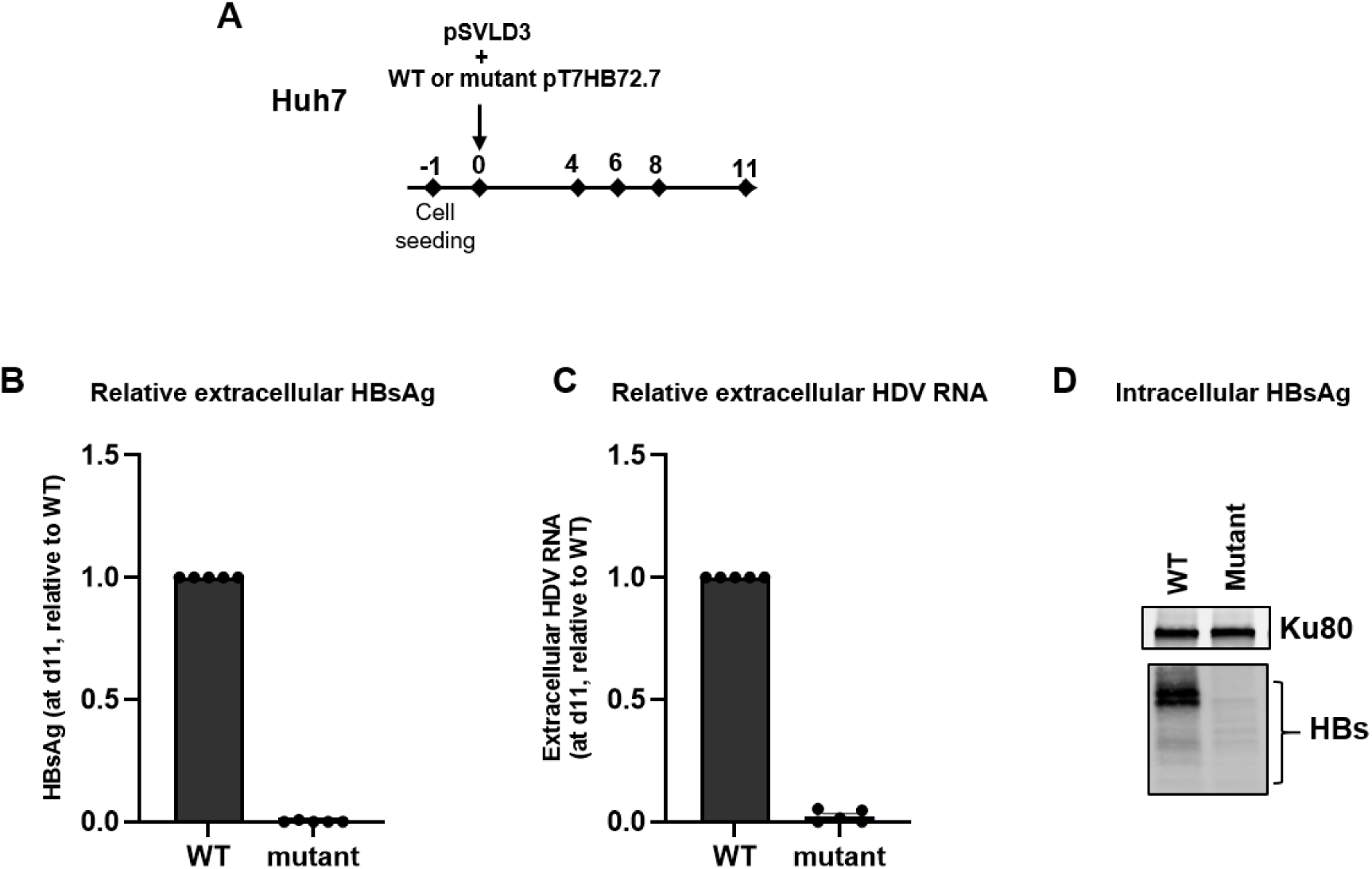
Adenine base editing of HBs ORF suppresses HDV release in the co-transfection model of pSVLD3 and pT7HB2.7 harboring the gS2 mutations. (A) Schematic of the protocol used to co-transfect Huh7 cells with pSVLD3 and WT or mutant pT7HB2.7. (B) Relative HBsAg (C) and extracellular HDV RNA levels were plotted for day 11 (d11). (D) Western blot to detect intracellular HBsAg. Ku80 served as a protein loading control. Data are represented as mean ± SEM.

Similar to the experiments with PLC/PRF/5 cells, a robust decrease in HBsAg was accompanied by the complete suppression of extracellular HDV RNA release (**Fig. 7B-7C**). The decrease of extracellular HBsAg was associated to reduced levels of intracellular HBsAg (**Fig. 7D**).

No differences in the levels of transfected plasmids or intracellular HDV RNA were observed in pSVLD3/WT-pT7HB2.7 vs pSVLD3/mutant-pT7HB2.7 transfected cells, confirming that the reduction in extracellular HDV RNA was not attributable to differences in transfection efficiency **(Fig. S5).**

When equal volumes of supernatants of PLC/PRF or Huh7 cells transfected with pSVLD3 were used to inoculate PHHs, a profound decrease in intracellular HDV RNA was observed in PHHs (**Fig. 8**) suggesting that the HBs editing mutations decreased significantly HDV release and further spread.

**Fig. 8.**
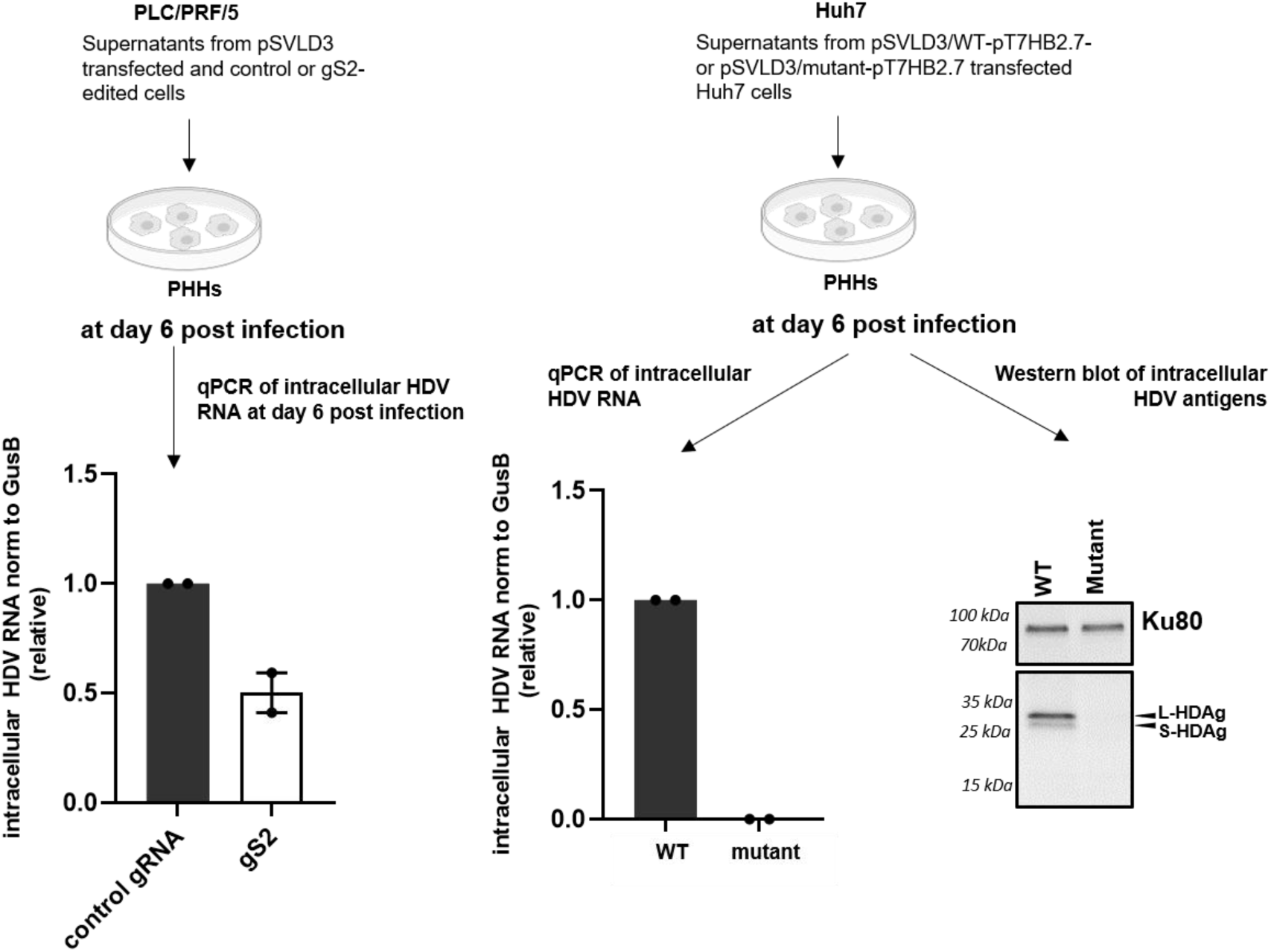
Adenine base editing of HBs ORF prevents HDV spread. PHHs were infected with equal volumes of supernatants collected from (A) PLC/PRF/5 cells which were first transfected with pSVLD3 followed by editing with either control gRNA or gS2 and (B) Huh7 cells co-transfected with pSVLD3/WT-pT7HB2.7 or pSVLD3/mutant-pT7HB2.7. Intracellular HDV RNA and protein levels were quantified on the 6th day post-infection.

## Discussion

Base editing, offering a method to introduce precise genetic changes, has revolutionized the field of biomedicine. With enhanced on-target and reduced off-target effects compared to CBEs, ABEs represent a significant advancement in the base editing tool kit. Through directed protein evolution, ABEs have progressed from the initially developed 7^th^-generation ABEs (48) to the highly efficient and therapeutically relevant 8^th^-generation of ABEs (ABE8s) (31,32). Recent studies have demonstrated the feasibility of liver-targeted delivery of ABEs, including ABE8s, highlighting their translational applicability to treat liver-associated diseases (43,49,50).

HBV is a DNA virus responsible for chronic viral infection associated with the persistence of both cccDNA and integrated sequences which represent ideal target for gene editing (13,46). In CHB patients, persistently elevated levels of HBsAg are associated with HBV specific B and T cell dysfunction, hindering immune-mediated control of HBV infection (51). A report by Michler T et al. suggested that restricting intracellular antigen expression is critical to break HBV immunotolerance (52). The current standard-of-care, NA therapy, effectively suppresses HBV replication but fails to reduce HBsAg levels. Approaches based on siRNA, ASO, and Cas13b can reduce HBsAg by targeting HBV transcripts. However, these agents do not directly target the HBV genomic reservoir, highlighting the need for new therapeutic approaches aiming at finite treatments. Here, we have investigated the potential of 8^th^ -generation ABEs to introduce mutations in the HBs ORF as a strategy to suppress HBsAg production. Our work demonstrated that this adenine base editing strategy can effectively inhibit HBsAg production from both the established pool of cccDNA and integrated HBV DNA by introducing mutations in the HBs ORF. Moreover, the use of mRNA-based delivery of ABEs via LNPs in vivo in HBV mouse models underscores its clinical translation potential.

Our in vitro cell culture experiments demonstrated that adenine base editing can robustly decrease HBsAg from HBV genotype D (HepG2-hNTCP, PHH, HepG2.2.15) or genotype A (PLC/PRF/5) models. Consistent with these findings, in vivo experiments using two mouse models-the HBVcircle mouse model (genotype D) and human liver chimeric mouse model (genotype C)- revealed sustained HBsAg reduction. Notably, the HBVcircle mouse model demonstrated up to 4-log decrease in HBsAg levels, whereas the human liver chimeric mouse model showed only a 1-log HBsAg reduction. While the two evaluated HBV-targeting gRNA recognize target sites that are conserved in the HBV genotypes in both mouse models, the observed differences in efficacy may arise from several other factors (53,54). Unlike the HBVcircle mouse model, the human liver chimeric mouse model allows reinfection of human hepatocytes, which may dilute the effects of base editing. Additionally, certain LNP formulations facilitate mRNA delivery more efficiently to mouse hepatocytes than human hepatocytes, potentially limiting the availability of LNP-encapsulated ABE mRNA/gRNA to human hepatocytes in chimeric models (55). It is also possible that the presence of anti-HBV immune response in the immunocompetent HBV-circle mouse model contributes to higher HBsAg reduction than the immunodeficient liver chimeric mouse model. Further work is needed to elucidate the precise mechanisms underlying the observed HBsAg suppression.

We also investigated whether adenine base editing of HBV could also have an antiviral effect on HDV infection. HDV requires the HBV envelope proteins for both the infection of hepatocytes and the release of infectious virus. It was shown that chronic HDV infection is sustained by intense HBsAg production from integrated HBV DNA in HBeAg-negative patients with very low levels of cccDNA (56,57). Globally, approximately 5% of CHB patients are co-infected with HDV with an increased risk of liver disease progression and complications (58). Despite the recent approval of Bulevirtide, an inhibitor of viral entry, in some parts of the world, new treatment options are still needed (11). Using PLC/PRF/5 cells, we showed in vitro evidence that suppressing HBsAg production from integrated HBV DNA by adenine base editing can be used as a strategy to inhibit HDV release and spread. These results open new avenues to investigate the effect of HBs base editing on HDV release in HBV/HDV coinfection in vitro in PHHs and in vivo in mouse models.

Taken together, our results suggest that, with its precision, efficiency, and translational potential, adenine base editing represents a promising new frontier for the treatment of CHB and CHD by reducing HBsAg and HBV replication, and inhibiting HDV release and spread.

## Acknowledgements

We acknowledge Inserm U1350 for discussions, Dr Birke Bartosch for providing the anti-HDAg antibody, Dr Christophe Combet and Dr David Durantel for the HBV genotype sequences A-H used in this study. We thank Dr Michel Rivoire from Centre Léon Bérard (Lyon, France) for providing human liver resections and the PHH platform staff of PaThLiv UMR U1350.

## Author Contributions

Conceptualization: AK, EC, EMS, MGM, GC, FG, MSP, BT, FZ; Formal Analysis: AK, EC, EMS, SD, DL, CYC, LM, MD, MSP; Funding acquisition: GC, FG, MSP, BT, FZ; Investigation: AK, EC, EMS, SD, DL, CYC, LM, MD, MLP, MSP; Methodology: AK, EC, EMS, EC, AK, DL, CYC, MLP, XG, MSP; Supervision: CS, FG, MSP, BT, FZ; Visualization: AK, EC, EMS, LM, MD, XG, BT; Writing – original draft: AK, EC, EMS, MSP, BT; Writing – review & editing: all authors.

## Footnotes

Author names in bold in references designate shared co-first authorship.

