## Supplementary Figures for "Adenine Base Editing Potently Suppresses Hepatitis B Surface Antigen Expression and Inhibits Hepatitis D Virus Release"

### Supplementary Fig. S1

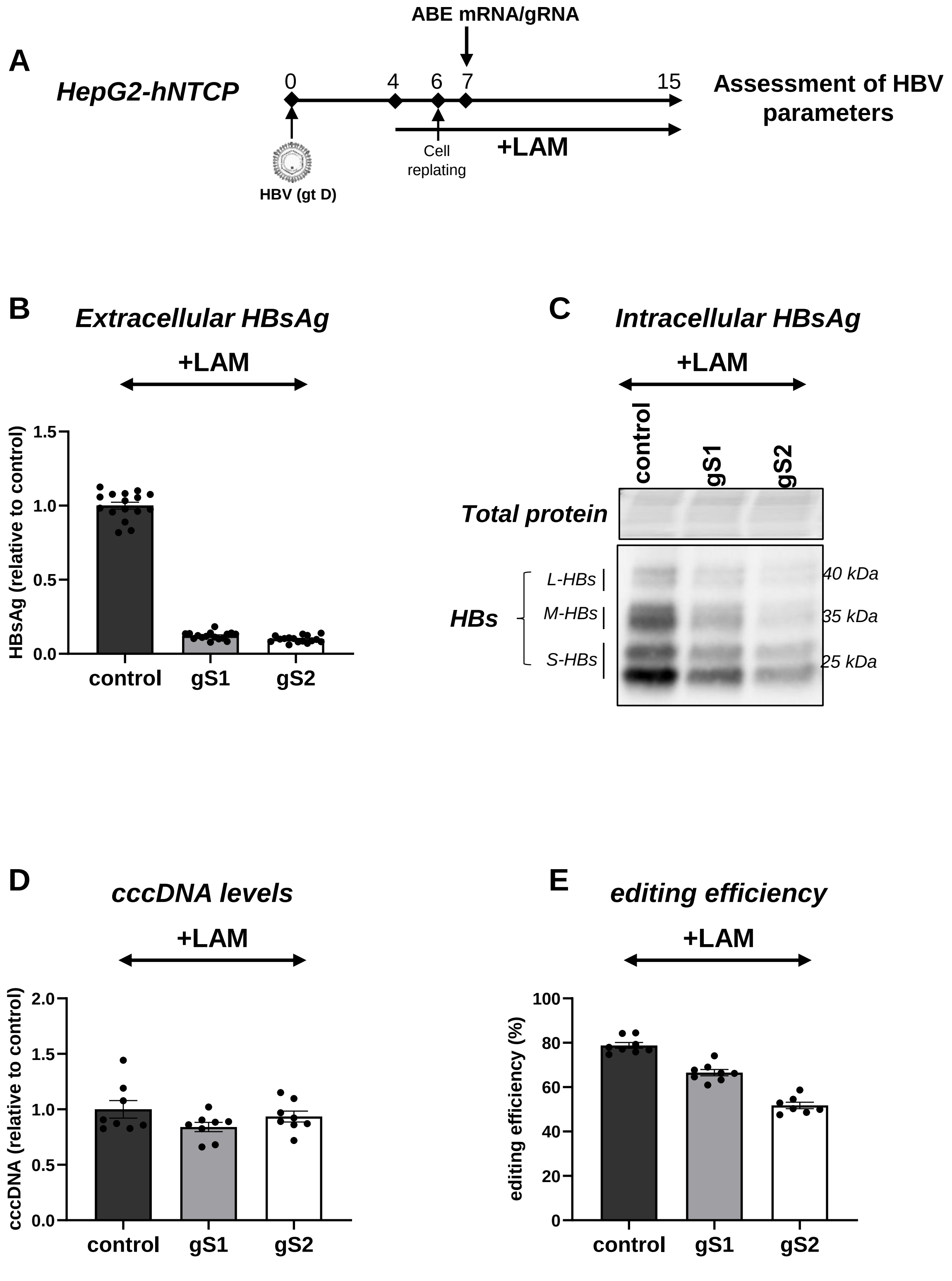


**Fig S1 Adenine base editing can be combined with NA.** (A) The schematic of the protocol used to transfect Lamivudine (LAM) pretreated HBV-infected HepG2-hNTCP cells with ABE-encoding mRNA and gS1 or gS2. (B-C) The levels of extracellular HBsAg and intracellular HBsAg were determined by ELISA and Western blot, respectively. (D) cccDNA levels were quantified by qPCR. (E) Percentage of A-to-G editing of HBs/POL ORF by gS1 and gS2 and control was determined on exonucleases I/III-treated samples and total extracted DNA samples, respectively. control: non-HBV targeting gRNA; LAM: Lamivudine. Data are represented as mean±SEM.

### Supplementary Fig. S2

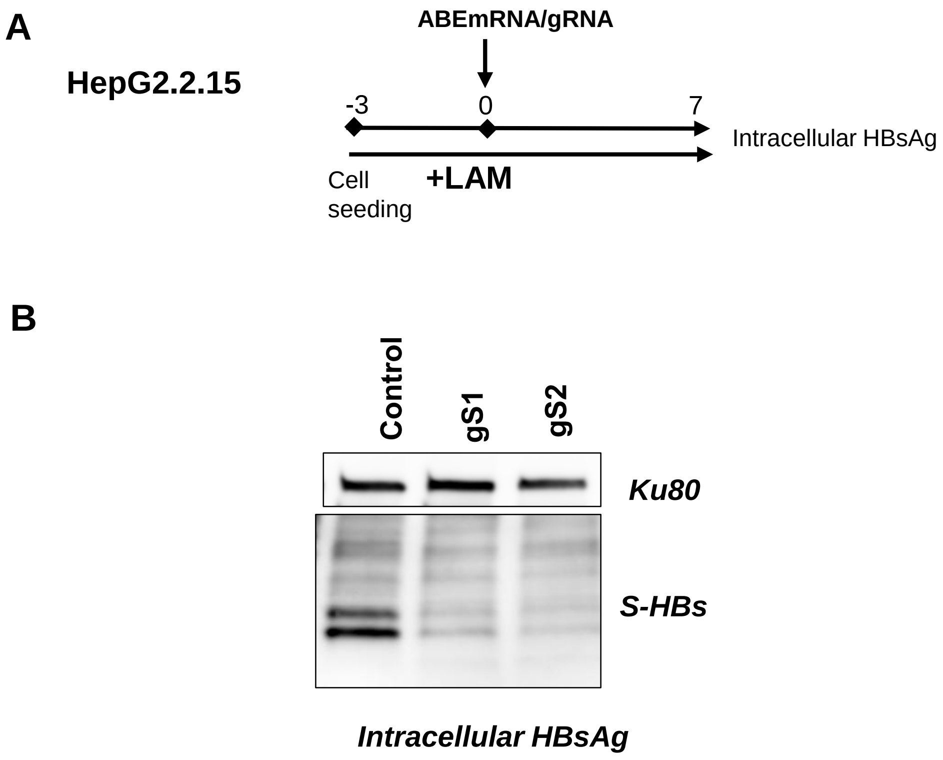


**Fig S2: ABE/gRNAs drastically reduce intracellular HBsAg from integrated HBV DNA** (A) The schematic of the protocol used to transfect LAM pretreated HepG2.2.15 cells with ABE-encoding mRNA and gS1 or gS2. (B) The levels of intracellular HBsAg was determined by Western blot. control gRNA: non-HBV targeting gRNA; LAM: Lamivudine

#### Supplementary Fig. S3

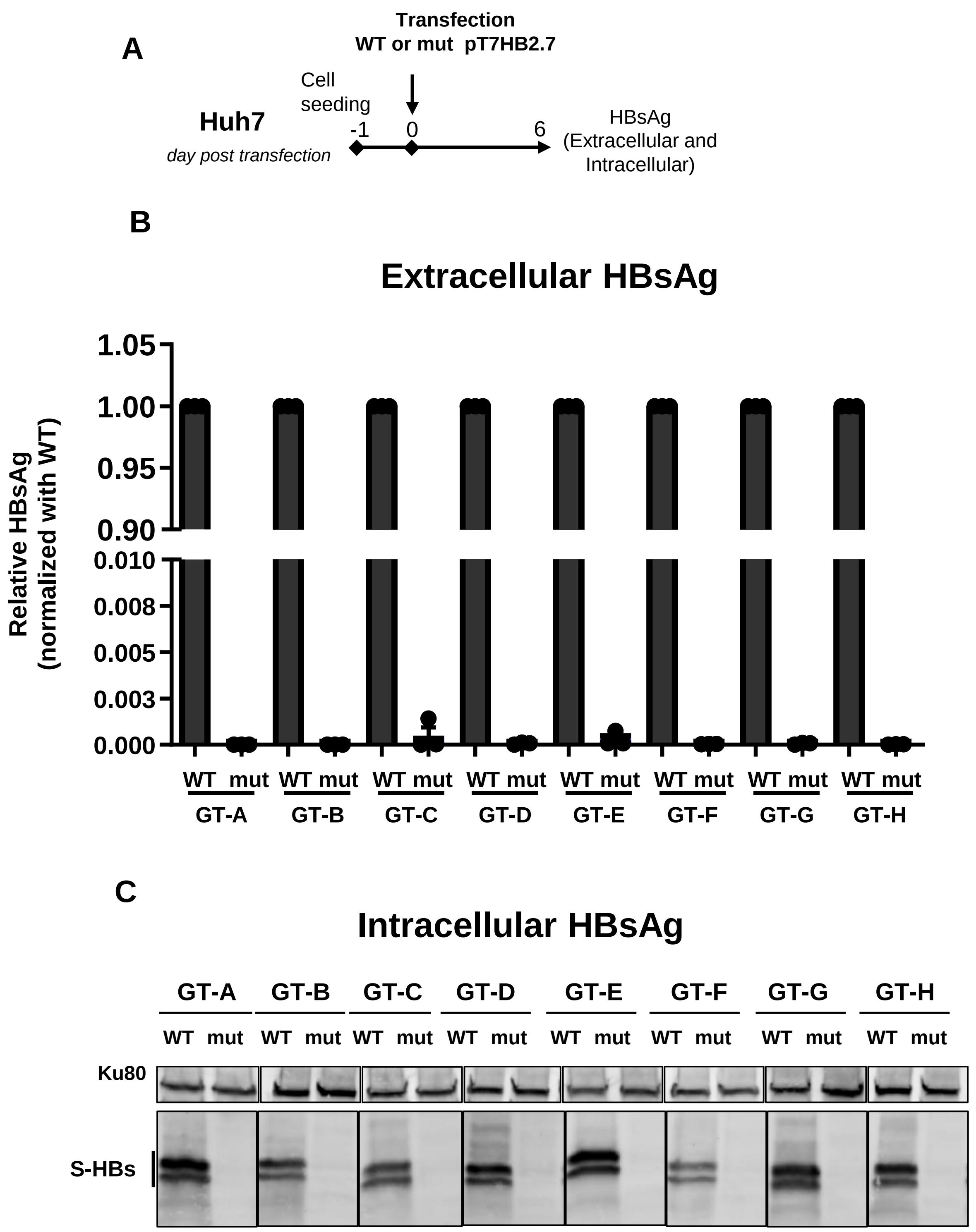


**Fig. S3**. (A) Experimental scheme (B-C) ELISA and Western blots showing the effect of gS2-corresponding mutations on secreted and intracellular HBsAg levels, respectively, across different HBV genotypes. Data were normalized with respect to WT condition for each genotype. (WT: wildtype; mut: mutant containing gS2-corresponding mutations; GT: HBV genotype; Letters A to H represent HBV genotypes)

#### Supplementary Fig. S4

A

PLC pSVLD3 levels

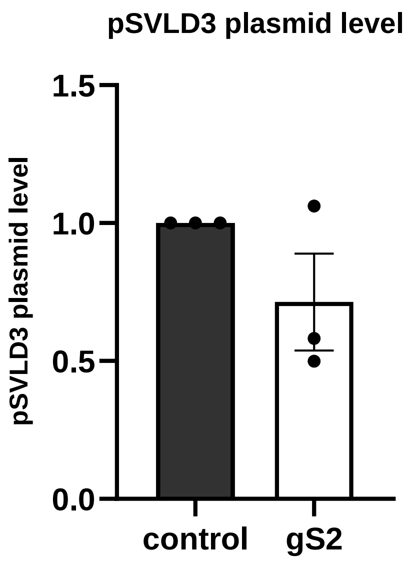


B

PLC/PRF/5 intracellular HDV RNA/pSVLD3

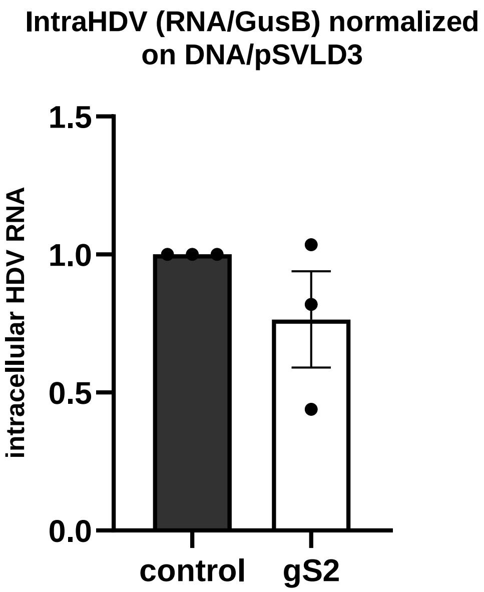


**Fig. S4**. For the experiment in Fig. 6, intracellular (A) pSVLD3 plasmid (B) HDV RNA levels were quantified from PLC/PRF/5 cells. The data were normalized with respect to control gRNA condition.

### Supplementary Fig. S5

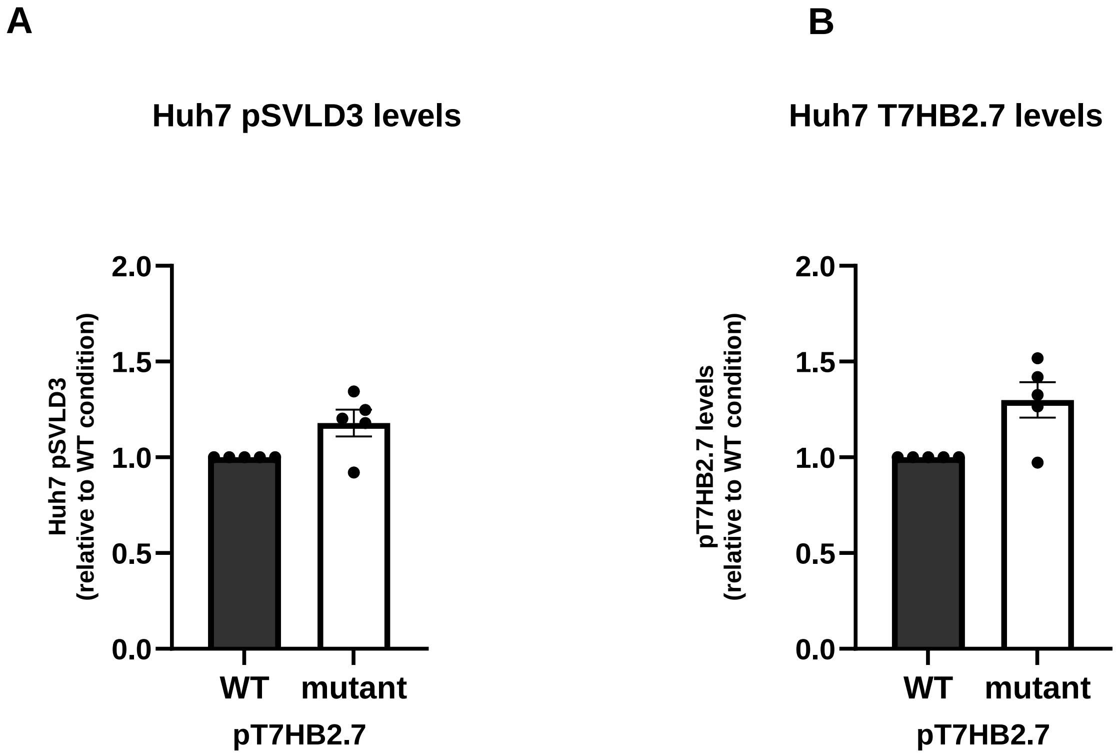


**Fig. S5**. For the experiment in Fig. 7, intracellular (A) pSVLD3 and (B) pT7HB2.7 plasmid levels were quantified from Huh7 cells. The data were normalized with respect to pT7HB2.7-WT condition.
